# Condensin dysfunction is a reproductive isolating barrier in mice

**DOI:** 10.1101/2022.10.20.513083

**Authors:** Warif El Yakoubi, Takashi Akera

**Affiliations:** Cell and Developmental Biology Center, National Heart, Lung, and Blood Institute, National Institutes of Health; Bethesda, Maryland 20894, USA

## Abstract

Reproductive isolation occurs when the genomes of two populations accumulate genetic incompatibilities that prevent inter-breeding. Cell biological understanding of such hybrid incompatibility is limited, especially for hybrid female sterility. Here we find that species divergence in condensin regulation and centromere organization between two mouse species, *Mus musculus* and *Mus spretus*, drives chromosome de-condensation and mis-segregation in their F1 hybrid oocytes, reducing female fertility. The chromosome condensation defects in hybrid oocytes were especially prominent at *Mus musculus* centromeres due to their highly abundant major satellite DNA, leading to species-specific chromosome mis-segregation. This study provides the first mechanistic insights into hybrid incompatibility in female meiosis and demonstrates that condensin mis-regulation can be a reproductive isolating barrier in mammals.

---

Hybrid incompatibility is the genetic cause of postzygotic reproductive isolation, significantly contributing to the speciation process (*1*). Such incompatibilities arise when the genomes of two closely related species or strains take distinct evolutionary trajectories, resulting in incompatible genomic interactions in F1 hybrids that lead to their lethality or sterility (*2*). Several hybrid incompatibility genes have been identified to explain hybrid lethality and hybrid male sterility (*3*–*6*). Although hybrid females exhibit fertility issues, facilitating reproductive isolation (*7, 8*), molecular mechanisms underlying hybrid female sterility are unknown.

Female meiosis I division is a highly error-prone process in many sexually reproducing organisms, affecting fertility (*9*–*12*). This error-prone nature implies that even the slightest form of hybrid incompatibility in this cell division can have a profound impact on reproduction. Female hybrid mice between *Mus musculus domesticus* (hereafter *musculus*) and *Mus spretus* (hereafter *spretus*) are sub-fertile due to chromosome segregation errors in meiosis I, creating aneuploid eggs (Fig. 1A, also see Materials and Methods, Mouse strains) (*13, 14*). Such meiotic failures would significantly contribute to maintaining reproductive isolation between these species. However, the molecular basis underlying hybrid incompatibility in the chromosome segregation process remained mysterious.

**Fig. 1.**
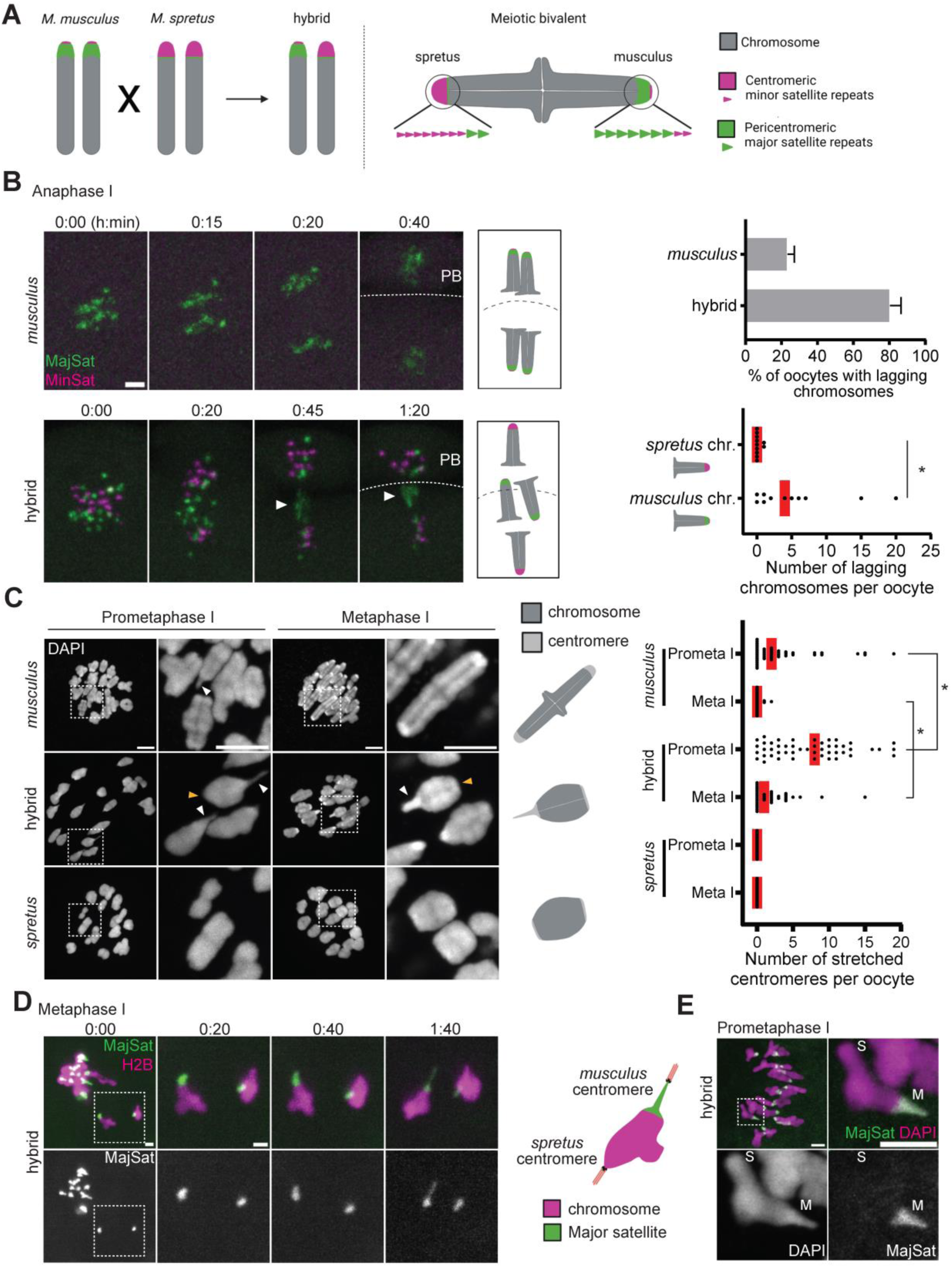
Species-specific chromosome mis-segregation driven by *musculus* centromere stretching. (**A**) Schematic of the hybrid mouse system to study hybrid incompatibility in female meiosis. (**B**) *musculus* and hybrid oocytes expressing TALE-mClover targeting major satellites (MajSat) and TALE-mRuby2 targeting *spretus* minor satellites (MinSat) were imaged live in anaphase I. MajSat and MinSat visualize *musculus* and *spretus* centromeres, respectively. White arrowheads, lagging chromosomes; PB, polar body; dashed lines, oocyte cortex. The anaphase lagging rate in *musculus* and hybrid oocytes (top graph) and the number of *spretus* and *musculus* chromosomes lagging in hybrid oocytes (bottom graph) were quantified (n = 20 oocytes for *musculus* and n = 11 for hybrid). (**C**) *musculus*, hybrid, and *spretus* oocytes were fixed at prometaphase I or metaphase I and counterstained with DAPI. White and orange arrowheads indicate stretched and compact centromeres, respectively. The number of stretched centromeres was quantified (n > 22 oocytes for each condition). (**D**) Hybrid oocytes expressing H2B-mCherry and MajSat were imaged live in metaphase I. (**E**) Hybrid oocytes expressing MajSat were fixed at prometaphase I. “M” and “S” indicate *musculus* and *spretus* centromeres, respectively. Images are maximum intensity z projections showing all chromosomes or optical slices magnified to show individual chromosomes. Each dot in the graph represents a single oocyte; red line, mean; *\*P* <0.05; scale bars, 5 *μ*m.

To understand the cell biological mechanism underlying the reduced fertility in *musculus* x *spretus* hybrid female, we imaged chromosome dynamics live in their oocytes. We found that chromosomes lagged in anaphase I more frequently in hybrid oocytes compared to control *musculus* oocytes, consistent with the higher aneuploid egg rate observed in this hybrid (Fig. 1B, top graph, and fig. S1A) (*13, 14*). Since centromere evolution can impact the chromosome segregation fidelity and centromere DNA have significantly diverged between *musculus* and *spretus*, we wonder if chromosomes from one parent could be more prone to segregation errors in this hybrid. Taking advantage of *spretus* centromeres having substantially more centromeric minor satellites and less pericentromeric major satellites compared to *musculus* centromeres (*15, 16*) (Fig. 1A), we can distinguish *musculus* and *spretus* centromeres using specific Transcription Activator-Like Effectors (TALE) constructs (*17, 18*). Differential labeling of parental chromosomes revealed that it was predominantly *musculus* chromosomes that were lagging in hybrid oocytes (Fig. 1B). To understand the mechanisms behind this species-specific chromosome mis-segregation, we analyzed their chromosome morphology in prometaphase I and metaphase I (Fig. 1C and fig. S1B to D). In *musculus* oocytes, sister chromatids were individualizing in prometaphase I with some centromeres still partially decondensed and stretched (Fig. 1C, *musculus*, white arrowhead). Chromosomes progressively condensed over time, and most centromeres were properly condensed by metaphase I in *musculus* oocytes. In contrast, hybrid oocytes had significantly decondensed chromosomes with centromeres stretched even in metaphase I (Fig. 1C and fig. S1B to D, hybrid). Centromere stretching was always observed just on one side of the meiotic bivalent in hybrid oocytes, implying a species-specific stretching of centromeres. Visualizing *musculus* centromeres with a major satellite TALE construct in both fixed and live cells revealed that it was major satellites at *musculus* centromeres that were stretching in hybrid oocytes (Fig. 1D and E). These results suggest that hybrid oocytes have chromosome condensation defects, especially impacting the *musculus* centromere structure, leading to their species-specific mis-segregation. This observation raises two fundamental questions: (i) why does hybrid oocytes have chromosome condensation defects, and (ii) why are major satellites most affected?

The condensin complex is a major factor to form the rigid chromosome structure in mitosis and meiosis (*19*). Since the chromosome condensation defects and centromere stretching in hybrid oocytes phenocopied the depletion of functional condensin II complex, the major condensin complex in mouse oocytes (*20*–*22*) (Fig. 2A), we hypothesized that condensin II is mis-regulated in hybrid oocytes. To test this hypothesis, we analyzed the localization of a condensin II-specific subunit, NCAPD3 (*20*), in hybrid oocytes and found that condensin II was significantly less enriched on the chromosome axis in hybrid oocytes compared to *musculus* oocytes (Fig. 2B and fig. S2A). Among hybrid oocytes, we noticed a variation in the condensin II abundance on the chromosome, which negatively correlated with the number of stretched centromeres per oocyte (fig. S2B). Moreover, reducing the condensin II abundance further in hybrid oocytes by partially depleting a condensin II-specific subunit, NCAPH2, enhanced centromere stretching (fig. S2C). These results collectively suggest that condensation defects and centromere stretching in hybrid oocytes are due to the reduced condensin II abundance on the chromosome.

**Fig. 2.**
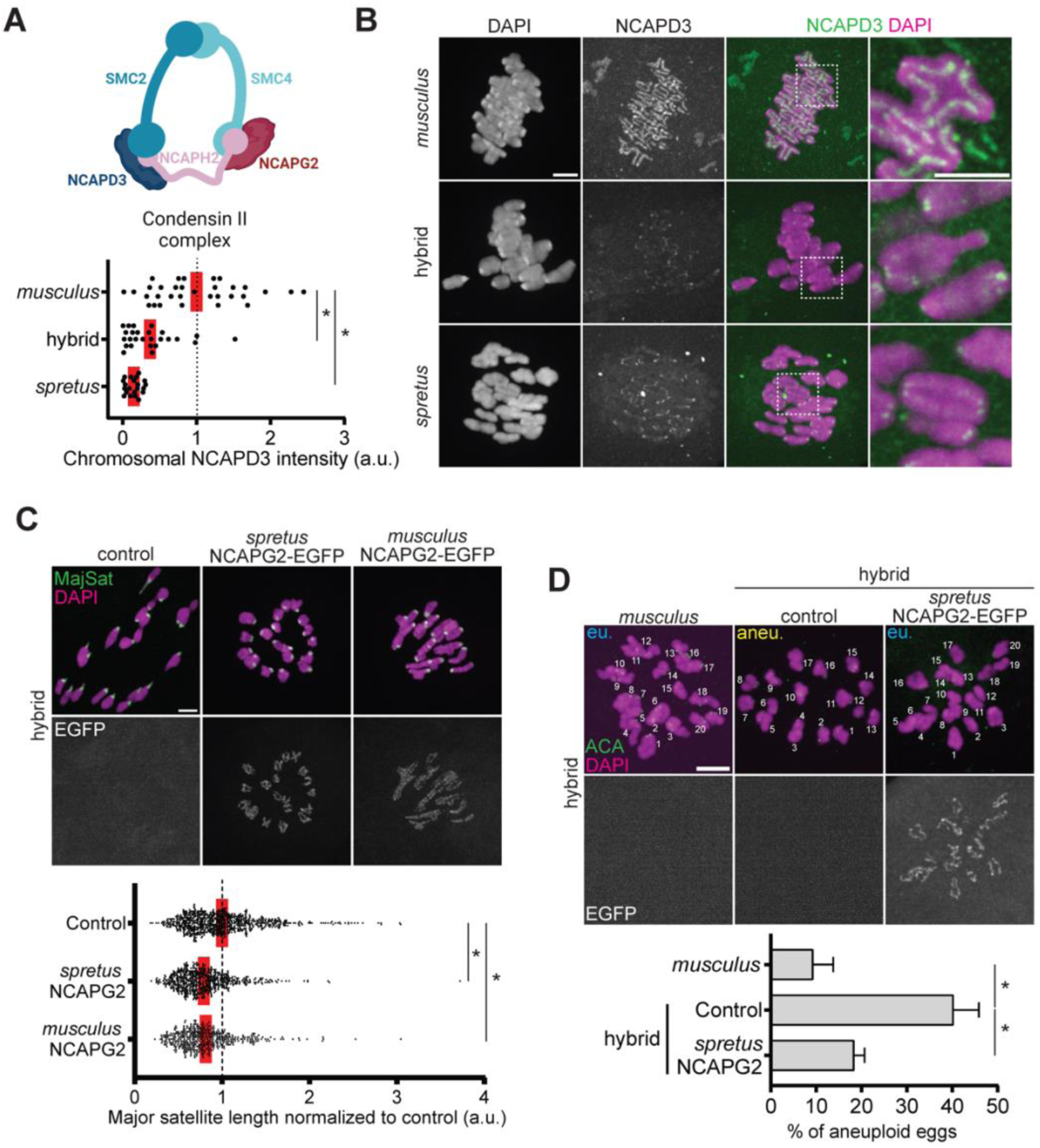
Condensin II dysfunction in hybrid oocytes causes egg aneuploidy. (**A**) Schematic of the condensin II complex. (**B**) *musculus*, hybrid, and *spretus* oocytes were fixed at metaphase I and stained for NCAPD3. NCAPD3 intensities on chromosomes were quantified (n > 21 oocytes for each condition). (**C**) Hybrid oocytes expressing NCAPG2^−EGFP^ derived from *musculus* or *spretus* were fixed at prometaphase I and stained for TOP2A (a major satellite marker, see fig. S7B) and EGFP. The length of major satellites was quantified; each dot in the graph represents a single centromere (n > 450 centromeres for each condition). (**D**) *musculus* oocytes and hybrid oocytes expressing *spretus* NCAPG2^−EGFP^ were matured to metaphase II, fixed, and stained for ACA (centromere) to perform the *in situ* chromosome counting assay. The aneuploid egg rate was quantified (n = 56, 66, and 21 cells for *musculus*, hybrid control, and hybrid + *spretus* NCAPG2, respectively); red line, mean; error bars, SD for three independent experiments. Images are maximum intensity z projections to show all chromosomes or optical slices to show individual chromosomes; *\*P* <0.05; scale bars: 5 *μ*m.

If the reduced condensin II levels on the chromosome is indeed the cause of centromere stretching in hybrid oocytes, overexpression of condensin II subunits may rescue this phenotype. To test this hypothesis, we overexpressed each of the condensin II-specific subunits, NCAPG2, NCAPD3, and NCAPH2 in hybrid oocytes (Fig. 2C and fig. S3). Among these subunits, the NCAPG2^−EGFP^ overexpression efficiently rescued centromere stretching (Fig. 2C). Importantly, overexpressing NCAPG2^−EGFP^ also rescued egg aneuploidy in the hybrid (Fig. 2D), demonstrating that condensin mis-regulation is one of the major hybrid incompatibilities affecting female fertility and maintaining the reproductive isolation between these species. Since *musculus* and *spretus* NCAPG2^−EGFP^ showed similar localization patterns and rescue efficiencies (Fig. 2C), their amino acid sequence divergence seems not to be part of the hybrid incompatibility in chromosome condensation (fig. S4).

Rescuing centromere stretching and aneuploidy by overexpressing a single condensin II subunit implies that NCAPG2 may be a rate-limiting factor to form a functional condensin II complex in this hybrid system. To test this idea, we examined the localization of condensin II subunits in late prophase I. Condensin II subunits are localized in the nucleus during interphase, providing them the opportunity to interact with chromatin prior to the entry into mitosis and meiosis (*20, 23*) (Fig. 3A and fig. S5A and B, *musculus*). We found that NCAPG2 was significantly less enriched in the nucleus in hybrid oocytes compared to *musculus* oocytes, while all other subunits enriched normally in the nucleus (Fig. 3A and fig. S5A and B). This specific reduction of NCAPG2 from hybrid oocyte nuclei can explain why NCAPG2 overexpression efficiently rescued centromere stretching and egg aneuploidy. Since total NCAPG2 protein levels were similar between *musculus* and hybrid oocytes (fig. S6), the nuclear-cytoplasmic transport of NCAPG2 could be differentially regulated between the two genotypes, reducing its nuclear localization specifically in hybrid oocytes.

**Fig. 3.**
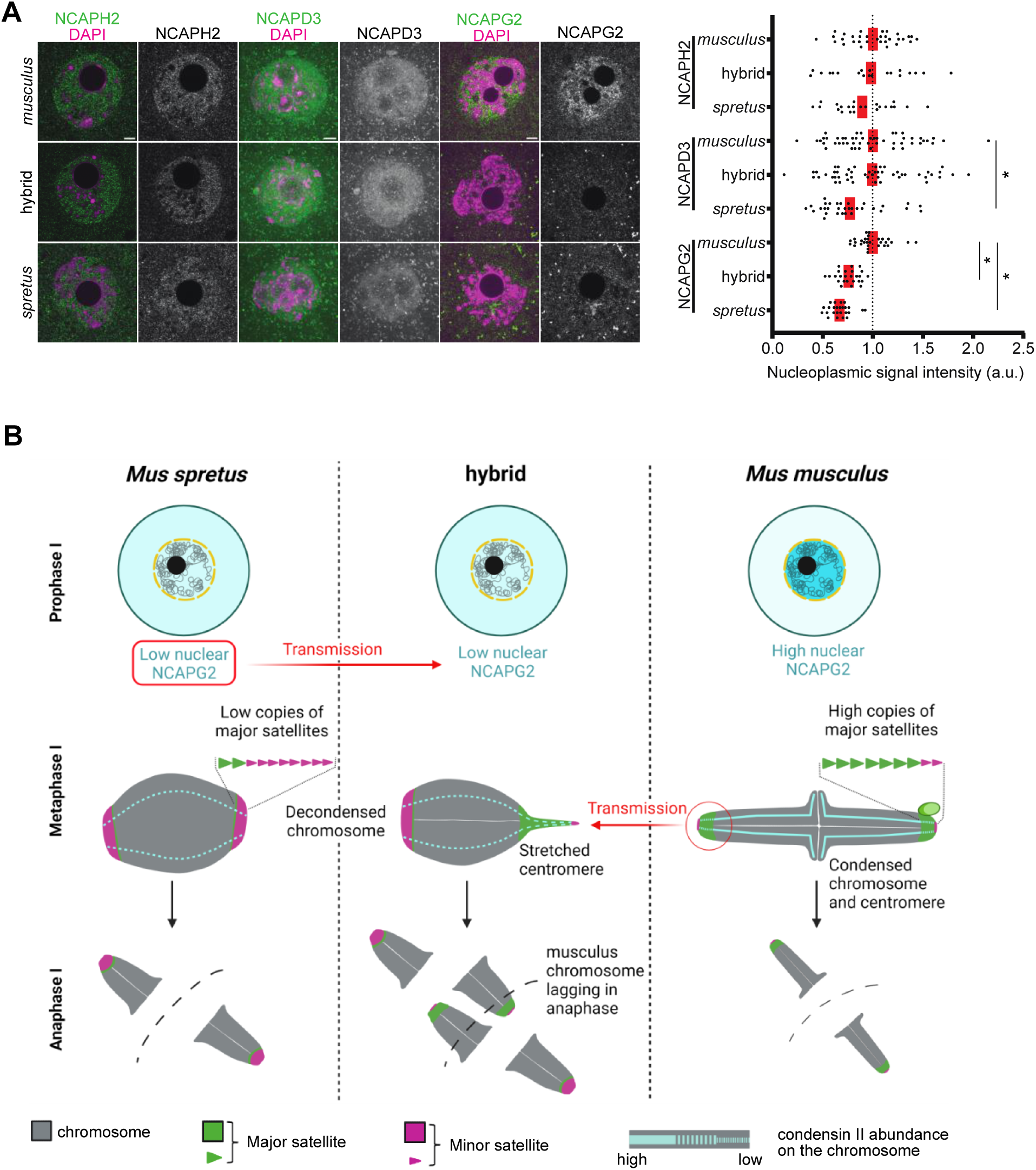
Species divergence in centromere organization and condensin regulation creates a reproductive isolating barrier in mice. (**A**) *musculus*, hybrid, and *spretus* oocytes were fixed at prophase I and stained for condensin II subunits. Their intensities in the nucleus were quantified; each dot in the graph represents a single oocyte (n > 20 oocytes for each condition); red line, mean. Nuclear NCAPG2 was reduced both in *spretus* and hybrid oocytes, while NCAPD3 was reduced in *spretus* but not in hybrid oocytes. Images are optical slices to show the nucleus; *\*P* <0.05; scale bars: 5 *μ*m. (**B**) Model of hybrid incompatibility reducing female fertility in *musculus* x *spretus* hybrid mice. Nuclear NCAPG2 levels are higher in *musculus* and lower in *spretus*, which dictates the abundance of condensin II on metaphase chromosomes. Chromosomes in *musculus* oocytes load higher levels of condensin II to condense major satellites, which is the chromosome region prone to stretching. In contrast, *spretus* oocytes enrich significantly less condensin II on the chromosome, but this low abundance is sufficient to condense their centromeres, which have very little major satellites. Hybrid oocytes inherited the low condensin II trait from *spretus* and the high major satellite copy number from *musculus*, and this combination drives centromere stretching, causing meiotic failures.

Interestingly, pure *spretus* oocytes also showed reduced NCAPG2 levels in prophase I nuclei and low condensin II levels on metaphase chromosomes, mimicking the localization patterns in hybrid oocytes (Fig. 2B and 3A and fig. S2A and S5B). This similarity in condensin dynamics between hybrid and *spretus* oocytes implies that hybrids inherited this reduced condensin II trait from *spretus*, eventually leading to stretching *musculus* centromeres in hybrid oocytes. Importantly, even though pure *spretus* oocytes have very low chromosomal condensin II levels, resulting in less compacted and less individualized chromosomes, their centromeres were not stretched (Fig. 1C and fig. S1B to D). This could be explained by *spretus* centromeres having very little major satellites, which is the chromosome region prone to stretching (Fig. 3B). In contrast, *musculus* oocytes enrich higher levels of condensin II, so that they have enough abundance to condense their major satellites. These results demonstrate a significant species divergence in the essential chromosome condensation process between *Mus musculus* and *Mus spretus*. As a result, hybrid oocytes inherit the reduced condensin II trait from *spretus* and major satellites from *musculus*, and we propose that this combination is what causing the major satellite stretching, leading to species-specific chromosome mis-segregation and reduced fertility.

Next, we asked why major satellites are prone to stretching. Analyzing the condensin II localization pattern within the chromosome revealed that major satellites enriched especially less condensin II compared to the rest of the chromosome, including the *spretus* centromere (*24*) (Fig. 4A). This localization pattern was also observed in *musculus* oocytes (fig. S7A), indicating that condensin II has an intrinsic property of loading less on major satellites. To understand why major satellites enrich less condensin II, we focused on Topoisomerase IIa (TOP2A), which is another major player in chromosome structure enriched at major satellites (*20, 25*) (fig. S7B). We found that the TOP2A abundance at major satellites correlates with the number of stretched centromeres in hybrid oocytes (Fig. 4B). This observation raises two possibilities: (i) TOP2A promotes centromere stretching by reducing condensin II abundance at major satellites or (ii) TOP2A is enriched at major satellites as a result of centromere stretching to fix the stretching. To distinguish between these models, we overexpressed TOP2A in hybrid oocytes (fig. S7C). The major satellite length was significantly longer when overexpressing ^EGFP-^TOP2A, showing that TOP2A enrichment promotes centromere stretching rather than rescuing it. To test if the centromeric enrichment of TOP2A is sufficient to induce the stretching by reducing condensin II abundance, we ectopically enriched TOP2A to *spretus* centromeres where TOP2A levels are usually very low (Fig. 4C and fig. S7B). To target TOP2A to *spretus* centromeres in hybrid oocytes, we fused TOP2A to a TALE construct specific to *spretus* minor satellites (MinSat-TOP2A) (*18*) (Fig. 4C). This targeting approach recruited TOP2A to *spretus* centromeres at a similar level to major satellites (Fig. 4C and fig. S8A). Importantly, the ectopic TOP2A targeting significantly reduced condensin II levels at *spretus* centromeres and induced their stretching (Fig. 4D and E). These results demonstrate that the centromeric TOP2A enrichment reduces condensin II abundance and induce centromere stretching in oocytes (Fig. 4F). Condensins drive DNA loop extrusion with their ATPase domains (*26*), and it is thought that they fall off from chromosomes when they cannot efficiently perform loop extrusion (*27, 28*). We propose that high TOP2A enrichment induces significant DNA inter-catenation at major satellites (*29*), which restricts condensin II-dependent loop extrusion, ultimately leading to a decrease in condensin II at this chromosome region. Supporting this idea, targeting TOP2A catalytic-dead mutant (TOP2A^Y804F^), which cannot induce DNA catenation, and a TOP2A mutant lacking the C-terminal domain (TOP2A^ΔCTD^), which cannot guide itself to the chromosome axis where condensin II localizes (*30*), reduced the condensin II abundance and stretched the centromere less efficiently compared to wild-type TOP2A (Fig. 4D and E and fig. S8B and C).

**Fig. 4.**
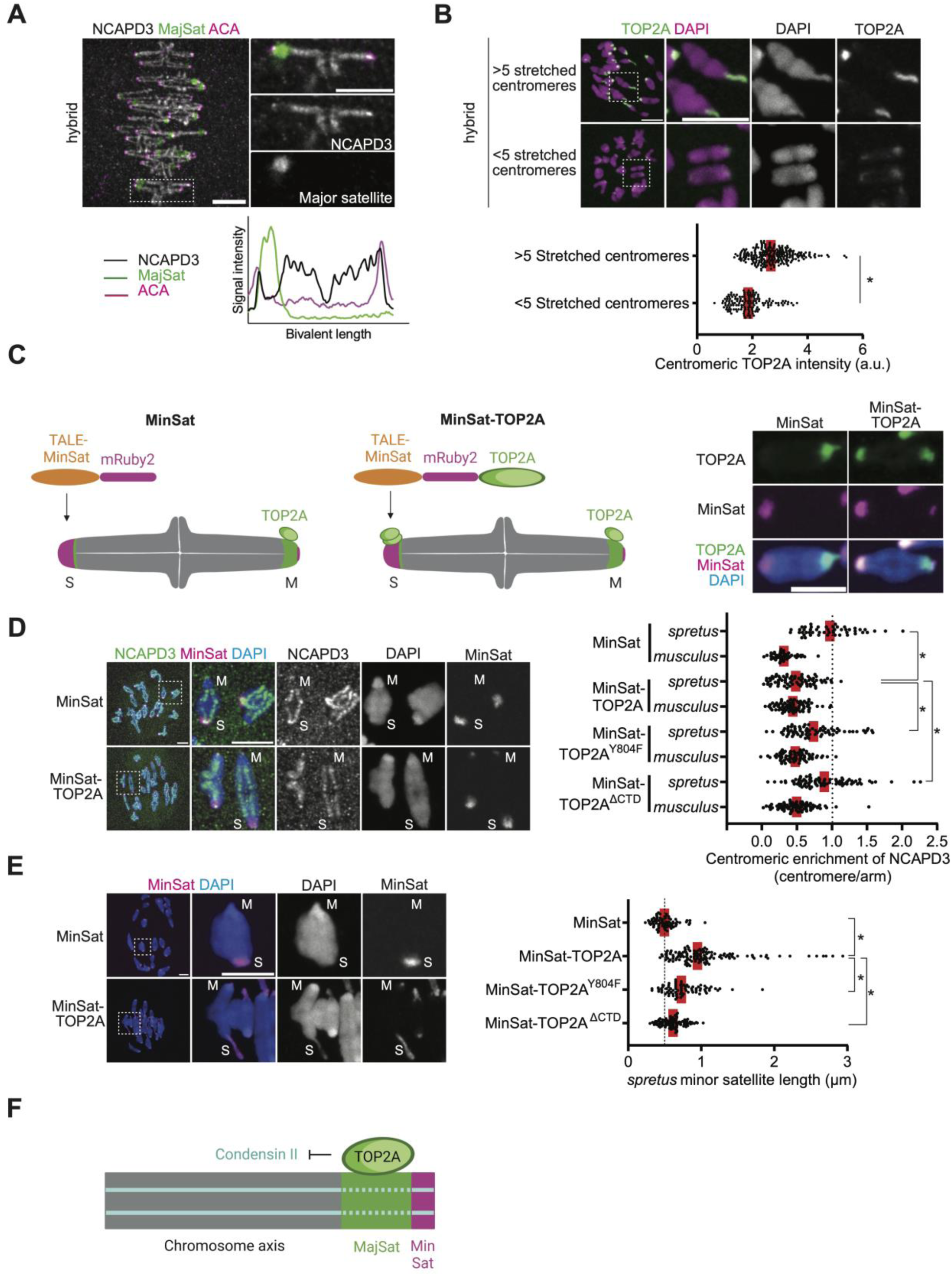
TOP2A reduces condensin II abundance at major satellites, promoting centromere stretching. (**A**) Hybrid oocytes expressing MajSat were fixed at metaphase I and stained for NCAPD3 and ACA (centromere). The graph is line scans of relative signal intensities across the chromosome length. (**B**) Hybrid oocytes were fixed at prometaphase I and stained for TOP2A. Oocytes were categorized into two groups based on the number of stretched centromeres, and their TOP2A signal intensities at *musculus* centromeres were quantified. (**C**) Schematic of the strategy to target TOP2A to *spretus* minor satellites. Hybrid oocytes expressing MinSat or MinSat-TOP2A were fixed at prometaphase I and stained for TOP2A. (**D, E**) Hybrid oocytes expressing MinSat, MinSat-TOP2A, MinSat-TOP2A^Y804F^, or MinSat-TOP2A^ΔCTD^ were fixed either at metaphase I and stained for NCAPD3 (D) or fixed at prometaphase I and counterstained with DAPI (E). Graphs show centromeric enrichment of NCAPD3, calculated as the centromeric signal divided by the chromosome arm signal for each half-bivalent (D) or the length of *spretus* minor satellites (E). Images are maximum intensity z projections to show all chromosomes or optical slices magnified to show individual chromosomes. “M” and “S” indicate *musculus* and *spretus* centromeres, respectively. Each dot in the graph represents a single centromere (n > 50 centromeres for each condition); red line, mean; *\*P* <0.001; scale bars: 5 *μ*m. (**F**) Schematic showing that high TOP2A enrichment at major satellites reduces the condensin II abundance.

This work provides the first cell biological insights into hybrid incompatibility in female meiosis, revealing that species divergence in centromere organization and condensin regulation drives meiotic failures, creating a reproductive isolating barrier (Fig. 3B). *musculus* x *spretus* hybrids also have male fertility issues mainly due to the defective chromosome pairing during meiotic prophase (*31*). Interestingly, a recent study showed that rescuing the pairing defects in male hybrids only partially restored their fertility, raising a possibility that other incompatibilities such as condensin dysfunction at play, further reducing their fertility (*32*). This work also suggest that the balance between condensin and TOP2A activities at the centromere is critical to properly segregate chromosomes, consistent with a previous study in *Drosophila* embryonic mitosis (*33*).

Condensin II is implicated in centromere drive, where selfish centromeres preferentially segregate to the egg to increase their own transmission rate (*24, 34*). Evolution of condensin regulations, including how they interact with centromere satellites, could be a strategy of the genome to fight back selfish centromeres to suppress their cheating in female meiosis (*35, 36*). Therefore, this reproductive isolating barrier could be a consequence of genetic conflict between selfish elements and the rest of the genome (*3, 37*).

## Acknowledgements

We thank A.E. Kelly, N.M. Rusan, and N. Phadnis for comments on the manuscript, M.A. Lampson, B.E. Black, and the Akera lab members for discussion, T. Hirano for the NCAPD3 antibody, and M.E. Torres-Padilla and Y. Miyanari for the TALE constructs. Schematics in the figures are created with BioRenders.com.

## Funding

Division of Intramural Research at the National Institutes of Health/National Heart, Lung, and Blood Institute, 1ZIAHL006249 (TA).

## Author contributions

Conceptualization: TA; Methodology: WE, TA; Investigation: WE, TA; Funding acquisition: TA; Supervision: TA; Writing – original draft: WE; Writing – review & editing: WE, TA.

## Competing interests

Authors declare that they have no competing interests.

## Supplementary Materials

Materials and Methods

figs. S1 to S8

References (38–41)

## Notes

### Competing Interest Statement

The authors have declared no competing interest.

## References

1. J. A. Coyne, H. A. Orr, Speciation (Sinauer, Sunderland, MA, 2004).

2. H. A. Orr, J. P. Masly, D. C. Presgraves, Speciation genes. Curr. Opin. Genet. Dev. 14, 675–9 (2004).

3. N. A. Johnson, Hybrid incompatibility genes: remnants of a genomic battlefield? Trends Genet. 26, 317–25 (2010).

4. O. Mihola, Z. Trachtulec, C. Vlcek, J. C. Schimenti, J. Forejt, A mouse speciation gene encodes a meiotic histone H3 methyltransferase. Science. 323, 373–5 (2009).

5. N. J. Brideau, H. A. Flores, J. Wang, S. Maheshwari, X. Wang, D. A. Barbash, Two Dobzhansky-Muller genes interact to cause hybrid lethality in Drosophila. Science. 314, 1292–5 (2006).

6. N. Phadnis, E. P. Baker, J. C. Cooper, K. A. Frizzell, E. Hsieh, A. F. A. de la Cruz, J. Shendure, J. O. Kitzman, H. S. Malik, An essential cell cycle regulation gene causes hybrid inviability in Drosophila. Science. 350, 1552–5 (2015).

7. T. A. Suzuki, M. W. Nachman, Speciation and reduced hybrid female fertility in house mice. Evolution. 69, 2468–81 (2015).

8. A. H. Sturtevant, Genetic Studies on DROSOPHILA SIMULANS. I. Introduction. Hybrids with DROSOPHILA MELANOGASTER. Genetics. 5, 488–500 (1920).

9. T. Chiang, R. M. Schultz, M. A. Lampson, Meiotic origins of maternal age-related aneuploidy. Biol. Reprod. 86, 1–7 (2012).

10. T. S. Kitajima, Mechanisms of kinetochore-microtubule attachment errors in mammalian oocytes. Dev. Growth Differ. 60, 33–43 (2018).

11. S. I. Nagaoka, T. J. Hassold, P. A. Hunt, Human aneuploidy: mechanisms and new insights into an age-old problem. Nat. Rev. Genet. 13, 493–504 (2012).

12. C. Thomas, T. Cavazza, M. Schuh, Aneuploidy in human eggs: contributions of the meiotic spindle. Biochem. Soc. Trans. 49, 107–118 (2021).

13. J. Sebestova, A. Danylevska, L. Novakova, M. Kubelka, M. Anger, Lack of response to unaligned chromosomes in mammalian female gametes. Cell Cycle. 11, 3011–8 (2012).

14. K. E. Koehler, S. E. Schrump, J. P. Cherry, T. J. Hassold, P. A. Hunt, Near-human aneuploidy levels in female mice with homeologous chromosomes. Curr. Biol. 16, R579–80 (2006).

15. S. Narayanswami, N. A. Doggett, L. M. Clark, C. E. Hildebrand, H. U. Weier, B. A. Hamkalo, Cytological and molecular characterization of centromeres in Mus domesticus and Mus spretus. Mamm. Genome. 2, 186–94 (1992).

16. A. K. C. Wong, F. G. Biddle, J. B. Rattner, The chromosomal distribution of the major and minor satellite is not conserved in the genus Mus. Chromosoma. 99, 190–195 (1990).

17. J. Reichmann, B. Nijmeijer, M. J. Hossain, M. Eguren, I. Schneider, A. Z. Politi, M. J. Roberti, L. Hufnagel, T. Hiiragi, J. Ellenberg, Dual-spindle formation in zygotes keeps parental genomes apart in early mammalian embryos. Science. 361, 189–193 (2018).

18. Y. Miyanari, C. Ziegler-Birling, M.-E. Torres-Padilla, Live visualization of chromatin dynamics with fluorescent TALEs. Nat. Struct. Mol. Biol. 20, 1321–1324 (2013).

19. T. Hirano, Condensin-Based Chromosome Organization from Bacteria to Vertebrates. Cell. 164, 847–57 (2016).

20. J. Lee, S. Ogushi, M. Saitou, T. Hirano, Condensins I and II are essential for construction of bivalent chromosomes in mouse oocytes. Mol. Biol. Cell. 22, 3465–77 (2011).

21. M. Houlard, J. Godwin, J. Metson, J. Lee, T. Hirano, K. Nasmyth, Condensin confers the longitudinal rigidity of chromosomes. Nat. Cell Biol. 17, 771–81 (2015).

22. M. M. Yoshida, K. Kinoshita, Y. Aizawa, S. Tane, D. Yamashita, K. Shintomi, T. Hirano, Molecular dissection of condensin II-mediated chromosome assembly using in vitro assays. Elife. 11, 1–27 (2022).

23. T. Ono, D. Yamashita, T. Hirano, Condensin II initiates sister chromatid resolution during S phase. J. Cell Biol. 200, 429–41 (2013).

24. T. Akera, E. Trimm, M. A. Lampson, Molecular Strategies of Meiotic Cheating by Selfish Centromeres. Cell. 178, 1132-1144.e10 (2019).

25. Y. Pommier, A. Nussenzweig, S. Takeda, C. Austin, Human topoisomerases and their roles in genome stability and organization. Nat. Rev. Mol. Cell Biol. 23, 407–427 (2022).

26. T. Terakawa, S. Bisht, J. M. Eeftens, C. Dekker, C. H. Haering, E. C. Greene, The condensin complex is a mechanochemical motor that translocates along DNA. Science. 358, 672–676 (2017).

27. M. Kong, E. E. Cutts, D. Pan, F. Beuron, T. Kaliyappan, C. Xue, E. P. Morris, A. Musacchio, A. Vannini, E. C. Greene, Human Condensin I and II Drive Extensive ATP-Dependent Compaction of Nucleosome-Bound DNA. Mol. Cell. 79, 99-114.e9 (2020).

28. K. Kinoshita, T. J. Kobayashi, T. Hirano, Balancing acts of two HEAT subunits of condensin I support dynamic assembly of chromosome axes. Dev. Cell. 33, 94–106 (2015).

29. T. Hsieh, Knotting of the circular duplex DNA by type II DNA topoisomerase from Drosophila melanogaster. J. Biol. Chem. 258, 8413–20 (1983).

30. K. Shintomi, T. Hirano, Guiding functions of the C-terminal domain of topoisomerase IIα advance mitotic chromosome assembly. Nat. Commun. 12, 2917 (2021).

31. D. W. Hale, L. L. Washburn, E. M. Eicher, Meiotic abnormalities in hybrid mice of the C57BL/6J x Mus spretus cross suggest a cytogenetic basis for Haldane’s rule of hybrid sterility. Cytogenet. Cell Genet. 63, 221–34 (1993).

32. B. Davies, A. Gupta Hinch, A. Cebrian-Serrano, S. Alghadban, P. W. Becker, D. Biggs, P. Hernandez-Pliego, C. Preece, D. Moralli, G. Zhang, S. Myers, P. Donnelly, Altering the Binding Properties of PRDM9 Partially Restores Fertility across the Species Boundary. Mol. Biol. Evol. 38, 5555–5562 (2021).

33. E. Piskadlo, A. Tavares, R. A. Oliveira, Metaphase chromosome structure is dynamically maintained by condensin I-directed DNA (de)catenation. Elife. 6, 1–22 (2017).

34. S. Henikoff, K. Ahmad, H. S. Malik, The centromere paradox: stable inheritance with rapidly evolving DNA. Science. 293, 1098–102 (2001).

35. T. D. King, C. J. Leonard, J. C. Cooper, S. Nguyen, E. F. Joyce, N. Phadnis, Recurrent Losses and Rapid Evolution of the Condensin II Complex in Insects. Mol. Biol. Evol. 36, 2195–2204 (2019).

36. C. Hoencamp, O. Dudchenko, A. M. O. Elbatsh, S. Brahmachari, J. A. Raaijmakers, T. van Schaik, Á. Sedeño Cacciatore, V. G. Contessoto, R. G. H. P. van Heesbeen, B. van den Broek, A. N. Mhaskar, H. Teunissen, B. G. St Hilaire, D. Weisz, A. D. Omer, M. Pham, Z. Colaric, Z. Yang, S. S. P. Rao, N. Mitra, C. Lui, W. Yao, R. Khan, L. L. Moroz, Kohn, J. St Leger, A. Mena, K. Holcroft, M. C. Gambetta, F. Lim, E. Farley, N. Stein, A. Haddad, D. Chauss, A. S. Mutlu, M. C. Wang, N. D. Young, E. Hildebrandt, H. H. Cheng, C. J. Knight, T. L. U. Burnham, K. A. Hovel, A. J. Beel, P.-J. Mattei, R. D. Kornberg, W. C. Warren, G. Cary, J. L. Gómez-Skarmeta, V. Hinman, K. Lindblad-Toh, F. Di Palma, K. Maeshima, A. S. Multani, S. Pathak, L. Nel-Themaat, R. R. Behringer, P. Kaur, R. H. Medema, B. van Steensel, E. de Wit, J. N. Onuchic, M. Di Pierro, E. Lieberman Aiden, B. D. Rowland, 3D genomics across the tree of life reveals condensin II as a determinant of architecture type. Science. 372, 984–989 (2021).

37. N. Phadnis, H. A. Orr, A single gene causes both male sterility and segregation distortion in Drosophila hybrids. Science. 323, 376–9 (2009).

38. A. Iwata-Otsubo, J. M. Dawicki-McKenna, T. Akera, S. J. Falk, L. Chmátal, K. Yang, B. Sullivan, R. M. Schultz, M. A. Lampson, B. E. Black, Expanded Satellite Repeats Amplify a Discrete CENP-A Nucleosome Assembly Site on Chromosomes that Drive in Female Meiosis. Curr. Biol. 27, 2365-2373.e8 (2017).

39. P. Stein, K. Schindler, Mouse oocyte microinjection, maturation and ploidy assessment. J. Vis. Exp. (2011), doi:10.3791/2851.

40. H. Igarashi, J. G. Knott, R. M. Schultz, C. J. Williams, Alterations of PLCbeta1 in mouse eggs change calcium oscillatory behavior following fertilization. Dev. Biol. 312, 321–30 (2007).

41. D. Clift, W. A. McEwan, L. I. Labzin, V. Konieczny, B. Mogessie, L. C. James, M. Schuh, A Method for the Acute and Rapid Degradation of Endogenous Proteins. Cell. 171, 1692-1706.e18 (2017).

