## Supplementary material for "Condensin dysfunction is a reproductive isolating barrier in mice": Material and Methods and Supplemental fig.

### Materials and Methods

Mouse strains

Mouse strains were purchased from Envigo (NSA, stock# 033 corresponds to CF1, *Mus musculus*), from Jackson Laboratory (C57BL/6J, stock# 000664, *Mus musculus*), and from RIKEN BioResource Research Center (SPR2, stock# RBRC00208, *Mus spretus*). CF-1 and C57BL/6J share the same centromere organization (*38*) and have complementary advantages as *Mus musculus* strains: CF-1 is an outbred strain with a significantly higher oocyte yield, and C57BL/6J is an inbred strain that efficiently produces hybrid offspring with SPR2. *Mus musculus* C57BL/6J females were crossed to *Mus spretus* SPR2 males to generate F1 hybrids, because the other direction does not produce offspring. All animal experiments were approved by the Animal Care and Use Committee (National Institutes of Health Animal Study Proposal#: H-0327) and were consistent with the National Institutes of Health guidelines.

Mouse oocyte collection and culture

Germinal vesicle (GV)-intact oocytes were harvested from 6- to 10-week-old female mice in the M2 media (Sigma, cat# M7167) supplemented with 5 µM milrinone (Sigma, cat# 475840) to prevent meiotic resumption. After the oocyte collection, oocytes were transferred to the M16 media (Millipore, cat# M7292) containing 5 µM milrinone covered with parafin oil (Nacalai, cat# NC1506764) and incubated at 37°C in a humidified atmosphere of 5% $CO_2$ in air. To induce meiotic resumption, milrinone was washed out, and oocytes that did not undergo the nuclear envelope breakdown (NEBD) within 1.5 h after the milrinone washout were removed from the culture. For the *in situ* chromosome counting assay, oocytes were matured for 13 h after NEBD and subsequently treated with 100 µM monastrol (Millipore, cat# 475879) in the organ culture dish (Falcon, cat# 353037) for 2 h 15 min prior to the fixation (*39*).

Plasmid construction and mutagenesis

Constructs to express NCAPG2, NCAPD3, and NCAPH2 were created by amplifying their coding sequences from *M. m. domesticus* or *M. spretus* cDNA library by PCR and subcloning them into the plasmid *In Vitro* Transcription (pIVT) vector with an N-terminal EGFP for NCAPD3 and a C-terminal EGFP for NCAPG2 and NCAPH2 (*40*). The following primers were used for the PCR amplifications: 5'-tcaagcttgcatgcctgcagatggaaaaacgtgaggcgttca-3' and 5'-tgctcaccattctagatggattcagaaattctcccaaagt-3' for NCAPG2, 5'-tgcatggaggatgtggaggtgcgctttgctcac-3' and 5'-ccattctagagtcgacaggctgggccatggatgg-3' for NCAPH2, 5'-gctgtacaaggtcgacatggcgctgcaggatcttg-3' and 5'-cggtacccggggatcctcagttggctgtcttcagggg-3' for NCAPD3. Constructs to target TOP2A to *spretus* minor satellites were created by amplifying the TOP2A coding sequence from *M. m. domesticus* cDNA library by PCR and subcloning it into the TALE-MinSat vector (Addgene, cat# 47879). The following primers were used for the PCR amplification of TOP2A: 5'-atggacgagctgtacaagatggagttgtcaccgctgca-3' and 5'-ggtgcccatatgtacatcagaagaggtcgtcatcg-3'. To create the catalytic-dead TOP2A mutant construct, the active site tyrosine 804 was mutated to phenylalanine (Y804F) using the QuikChange Lightning Site-Directed Mutagenesis Kit (Agilent Technologies, cat # 210518) with the following primers: 5'-actcagctagtcctaggttcatctttacaatgctcag-3' and 5'-ctgagcattgtaaagatgaacctaggactagctgagt-3'. To create the TOP2A construct lacking the C-terminal domain (1-1190 aa), the MinSat-TOP2A plasmid was digested with Pfo I to remove the C-terminal domain (1191-1528 aa), and the digested plasmid was used as a DNA template to synthesize the *MinSat-Top2a*$^{\Delta CTD}$ cRNA (see below).

Oocyte microinjection

GV-intact oocytes were microinjected with ~5 pl of cRNAs or antibodies in M2 containing 5 µM milrinone, using a micromanipulator TransferMan 4r and FemtoJet 4i (Eppendorf). Following the microinjection, oocytes were maintained at prophase I in M16 supplemented with 5 µM milrinone overnight to allow protein expression. cRNAs used for microinjections were *Ncapg2-Egfp* (*M. musculus* or *M. spretus* NCAPG2 with EGFP at the C-terminus) at 700 ng/µl, *Egfp-Ncapd3* (*M. musculus* or *M. spretus* NCAPD3 with EGFP at the N-terminus) at 700 ng/µl, *Ncaph2-Egfp* (*M. musculus* or *M. spretus* NCAPH2 with EGFP at the C-terminus) at 700 ng/µl, *Egfp-Top2a* (*M. musculus* TOP2A fused to EGFP and LacI at the N-terminus) at 430 ng/µl, *MinSat-Top2a* (TALE construct that recognizes *M. spretus* minor satellite repeats fused to mRuby2 and *M. musculus* TOP2A at the C-terminus) at 1000 ng/µl, *MinSat-Top2a*$^{\Delta CTD}$ (TALE construct that recognizes *M. spretus* minor satellite repeats fused to mRuby2 and *M. musculus* TOP2A (aa 1-1190) at the C-terminus) at 1016 ng/µl, *MinSat-Top2a*$^{Y804F}$ (TALE construct that recognizes *M. spretus* minor satellite repeats fused to mRuby2 and *M. musculus* catalytic-dead TOP2A$^{\text{Y804F}}$ mutant at the C-terminus) at 1300 ng/µl, *MajSat* (TALE construct that recognizes major satellite repeats fused to mClover and 3 tandem Halo tag at the C terminus) at 1500 ng/µl, and *MinSat* (TALE construct that recognizes *M. spretus* minor satellite repeats fused to mRuby2 at the C-terminus; Addgene, cat# 47879) at 600 ng/µl, and *H2B-mCherry* (human histone H2B with mCherry at the C terminus) at 100 ng/µl. To TrimAway NCAPH2, *mCherry-Trim21* cRNA (*M. musculus* Trim21 fused with mCherry at the C-terminus, Addgene cat# 105522) at 800 ng/µl and rabbit anti-NCAPH2 antibody at 0.2 mg/ml (Invitrogen, cat# pa5-66964) were co-microinjected (*41*). cRNAs were synthesized using the T7 mMessage mMachine Kit (Ambion, cat# AM1340) and purified using the MEGAclear Kit (Thermo Fisher, cat# AM1908).

Whole oocyte and chromosome spread immunostaining

Oocytes and eggs were fixed at prometaphase I (3 h from the nuclear envelope breakdown) or at metaphase I (7 h from the nuclear envelope breakdown) in freshly prepared 2% paraformaldehyde (Electron Microscopy Sciences, cat# 15710) in 1x PBS (Quality Biological, car# 119-069-101CS) with 0.1% Triton X-100 (Millipore, cat# TX1568-1) for 20 min at room temperature (RT), permeabilized in 1x PBS with 0.1% Triton X-100 for 15 min at RT, placed in the blocking solution (0.3% BSA (Fisher bioreagents, cat# BP1600-100) and 0.01% Tween-20 (ThermoFisher, cat# J20605-AP) in 1x PBS) overnight at 4°C, incubated 2 h with primary antibodies at RT, washed three times for 10 min with the blocking solution, incubated 1 h with secondary antibodies at RT, washed three times for 10 min in the blocking solution, and mounted on microscope slides with the Antifade Mounting Medium with DAPI (Vector Laboratories, cat# H-1200). For the metaphase I NCAPD3 immunostaining, zona pellucida was removed with Acidic Tyrode's Solution (Millipore, cat# MR-004-D) 6 h after NEBD, incubated in M16 for 1 h, and fixed with 2% paraformaldehyde in 1x PBS with 0.3% Triton X-100 for 30 min at 37°C, incubated in the blocking / permeabilization solution (3% BSA in 1x PBS with 0.03% Triton X-100) overnight at 4°C. For chromosome spreads, zona pellucida was removed from oocytes microinjected with *MajSat* cRNA, and the oocytes were fixed with 1% paraformaldehyde, 0.15% Triton X-100, and 3 mM DTT (Sigma, cat# 43815) 7 h after NEBD. The following primary antibodies were used: rabbit anti-mouse NCAPD3 antibody (Fig. 2B, 3A, 4A, 4D, S2B, S7A,

S8B, 1:500, gift from Dr. T. Hirano), rabbit anti-human NCAPD3 (fig. S2A, 1:100, Bethyl, cat# A300-604A), rabbit anti-human NCAPG2 (Fig. 3A, 1:100; Bioss, cat# BS-7721R), rabbit anti-human NCAPG2 (fig. S5B, 1:100, Bethyl, cat# A300-605A), rabbit anti-human NCAPH2 (1:100, Bethyl, cat# A302-275A), goat anti-human CAPE (SMC2) (1:100, Abcepta, cat# AF2276a), rabbit anti-mouse SMC4 (1:100, Novus, cat# NBP1-86635), CREST human autoantibody against centromere (1:100, Immunovision, cat# HCT-0100), rabbit anti-Topoisomerase II (1:100, Abcam, cat# ab109524), goat anti-GFP antibody conjugated with Dylight488 (1:100, Rockland, cat# 600-141-215). Secondary antibodies were Alexa Fluor 488–conjugated donkey anti-rabbit (1:500, Invitrogen, cat# A21206) or donkey anti-goat (1:500, Invitrogen, cat# A11057), Alexa Fluor 568–conjugated goat anti-rabbit (1:500, Invitrogen, cat# A10042), or Alexa Fluor 647–conjugated goat anti-human (1:500, Invitrogen, cat# A21445).

Confocal microscopy

Fixed oocytes and eggs were imaged with a microscope (Eclipse Ti; Nikon) equipped with 100x 1.40 NA oil-immersion objective lens, CSU-W1 spinning disk confocal scanner (Yokogawa), ORCA Fusion Digital CMOS camera (Hamamatsu Photonics), and 405, 488, 561 and 640 nm laser lines controlled by the NIS-Elements imaging software (Nikon). Confocal images were acquired as z-stacks at 0.3 µm intervals. For live imaging, oocytes were placed into 3 µl drops of M2 covered with parafin oil in a glass-bottom tissue culture dish (fluoroDish, cat# FD35-100) in a stage top incubator (Tokai Hit) to maintain 37°C. Time-lapse images were collected with a microscope (Eclipse Ti2-E; Nikon) equipped with the 20x / 0.75 NA objective (Fig. 1B and fig. S1A) and 60x / 1.40 NA oil-immersion objective (Fig. 1D), CSU-W1 spinning disk confocal scanner (Yokogawa), ORCA Fusion Digital CMOS camera (Hamamatsu Photonics), and 405, 488, 561 and 640 nm laser lines controlled by the NIS-Elements imaging software (Nikon). Confocal images were collected as z-stacks at 1 µm intervals to visualize all the chromosomes. Images are displayed as maximum intensity z-projections.

Image analysis

Fiji/ImageJ (NIH) was used to analyze all the images. In general, optical slices containing chromosomes were added to produce a sum projection for pixel intensity quantifications. To quantify centromere signal intensities, ellipses were drawn around the centromeres, and signal intensity was integrated over each ellipse after subtracting background, obtained near the centromeres. To quantify chromosomal NCAPD3 signal intensities, masking images were created using DAPI staining images to specifically measure NCAPD3 signal intensities on the chromosome. For the *in situ* chromosome counting assay, the number of chromosomes were counted in metaphase II eggs using the DAPI and ACA (centromere) signals.

Western blot

Each oocyte sample contained 100 oocyte cells, and each thymus sample contained 5 mg of the thymus tissue. Thymus tissues were homogenized using the RIPA buffer (150 mM NaCl (Millipore, cat# 116224), 0.5% Sodium deoxycholate (Biochemica, cat# A1531) and 50 mM Tris, pH 8.0 (KD Medical, cat# RGF-3360), and 1% Triton X-100) supplemented with the protease inhibitor cocktail (MedChem Express, cat# HY-K0010). Oocyte samples and thymus cell extracts were mixed with Laemeli buffer (BioRad, cat# 1610737) containing β-mercaptoethanol (Sigma, cat# M6250) and denatured at 95°C for 5 min. Samples were separated on 10% Mini-Protean TGX Precast Protein Gels (Bio-Rad, cat# 4561036) or on 10% Mini-

Protean TGX Stain-Free Protein Gels (Bio-Rad, cat# 4568036) to visualize the total proteins with the GelDoc Go Gel Imaging System (Bio-Rad, cat# 12009077) before the transfer. Proteins were, transferred to the PVDF membrane (Bio-Rad, cat# 1704156) with the Trans-Blot Turbo Transfer System (Bio-Rad), and blocked for 1 h at RT with 2% ECL Prime Blocking agent (Cytiva, cat# RPN418V). The membrane was then incubated overnight at 4°C with the rabbit anti-human NCAPG2 (1:300, Bioss, cat# BS-7721R) followed by washing three times for 15 min each with the TBST buffer (20 mM Tris, pH 7.6, 150 mM NaCl, and 0.1% Tween-20) and incubation for 1 h with the rabbit peroxidase-conjugated secondary antibody (Bethyl, cat# A120-201P), which was detected using the ECL detection reagents (Bio-Rad, cat# 1705061) and Amersham Imager 600 (GE Healthcare).

Quantification and statistical analysis

Data points were pooled from at least two independent experiments (biological replicates) unless specified in the figure legend. Data analysis was performed using Microsoft Excel and GraphPad Prism 9. Column bar graphs and line and scattered plots were created with GraphPad Prism 9. Unpaired two-tailed t-test was used for statistical analysis, and a value of $P < 0.05$ was considered significant.

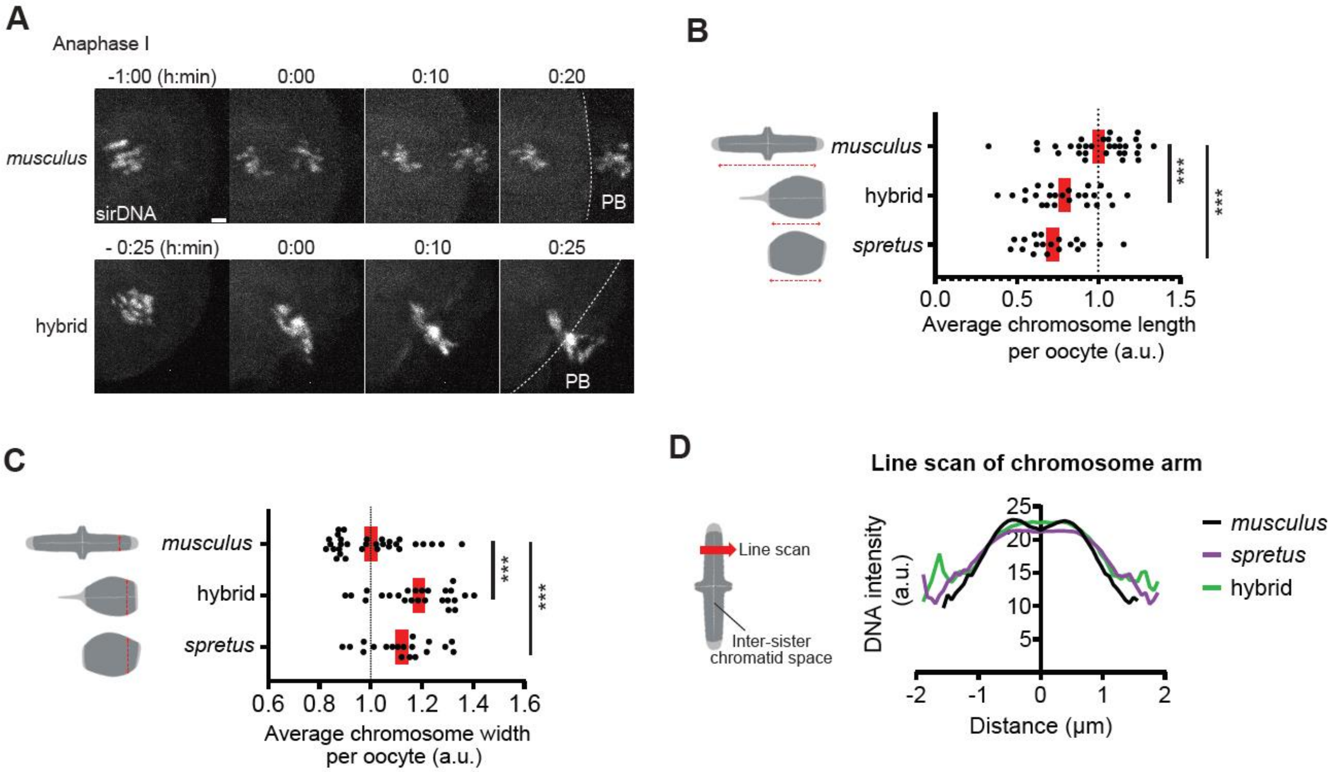


**Fig. S1.**
**Less condensed chromosomes in hybrid and *spretus* oocytes.** (**A**) *musculus* and hybrid oocytes were imaged live in anaphase I in the presence of sirDNA to visualize chromosomes. PB, polar body; dashed lines, oocyte cortex. See Fig. 1B for the quantification of lagging chromosomes. (**B**, **C**) The chromosome length (B) and width (C) were quantified. Each dot in the graph represents a single oocyte (n > 19 oocytes for each condition). (**D**) The graph shows line scans of the DNA intensity across the chromosome arm region to quantify the individualization of sister chromatids. *musculus* chromosomes are more individualized, indicated by the dip in the DNA intensity in the middle of the line scan (i.e., inter-sister chromatid space), compared to hybrid and *spretus* chromosomes. The line scans are averaged over n > 64 chromosomes for each genotype; red line, mean; ****P* <0.0001; scale bar: 5 $\mu$m.

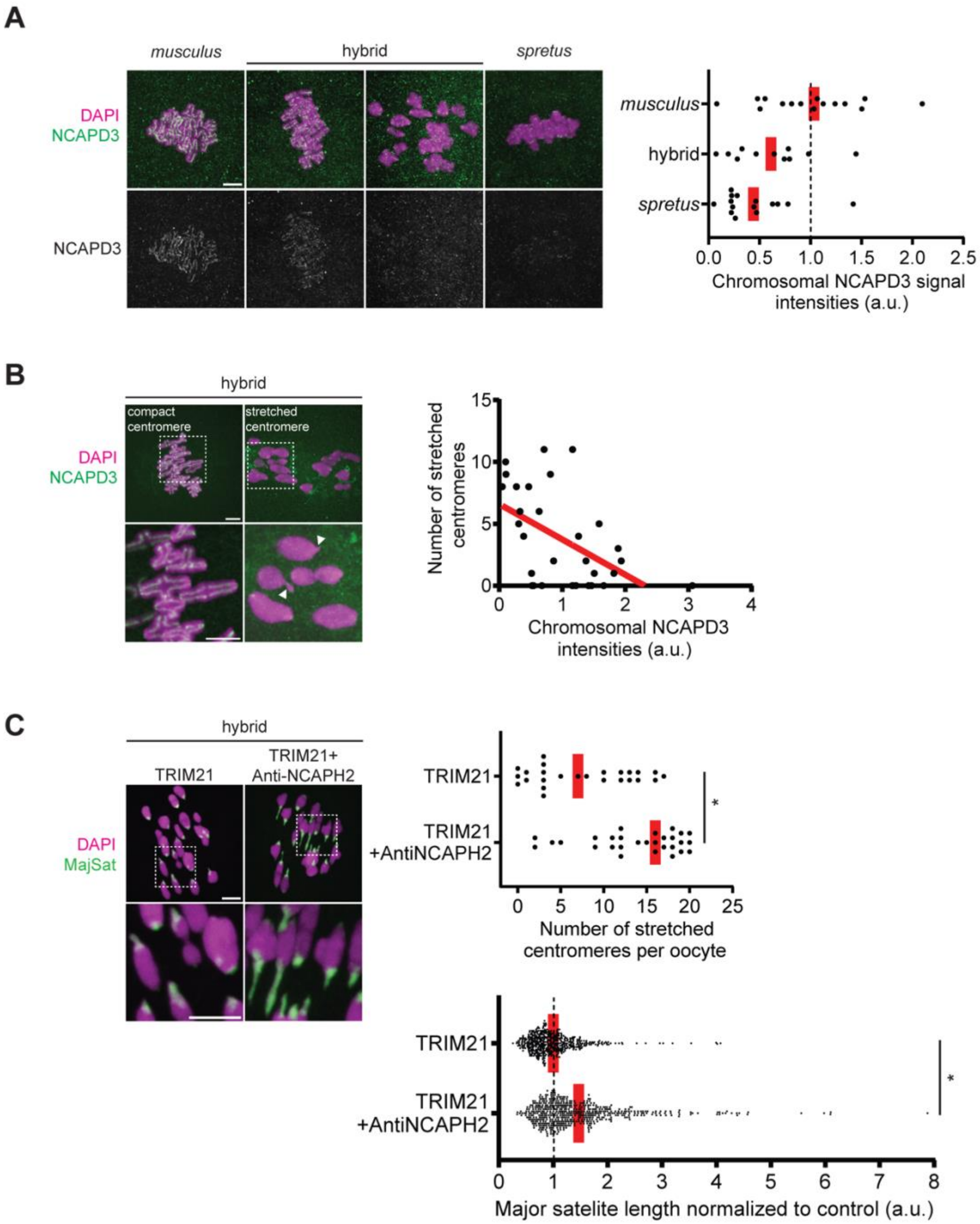


**Fig. S2.**

**Reduction in condensin II abundance leads to centromere stretching in hybrid oocytes.** (**A**) *musculus*, hybrid, and *spretus* oocytes were fixed at metaphase I and stained for NCAPD3, using a different antibody (Bethyl, cat# A300-604A) from Fig. 2B where the NCAPD3 antibody from the Hirano lab was used. NCAPD3 intensities on the chromosome were quantified, showing a similar trend with Fig. 2B (n > 11 oocytes for each condition). (**B**) Hybrid oocytes were fixed at metaphase I and stained for NCAPD3 (antibody from the Hirano lab). The chromosomal NCAPD3 intensities relative to the number of stretched centromeres per oocyte were plotted in the graph (n = 22 oocytes); white arrowheads, stretched centromeres; red line, a simple linear regression between the chromosomal NCAPD3 intensities and the number of stretched centromeres. (**C**) To partially deplete NCAPH2 by the TrimAway method (*41*), hybrid oocytes expressing mCherry-Trim21 with or without the anti-NCAPH2 antibody microinjection were fixed at prometaphase I and stained with TOP2A (a major satellite marker, see fig. S7B). The number of stretched centromeres per oocyte and the major satellite length were quantified (n > 25 oocytes for each condition). Images are maximum intensity z projections to show all chromosomes or optical slices magnified to show individual chromosomes; red line, mean; $*P < 0.05$; scale bars: 5 $\mu$m.

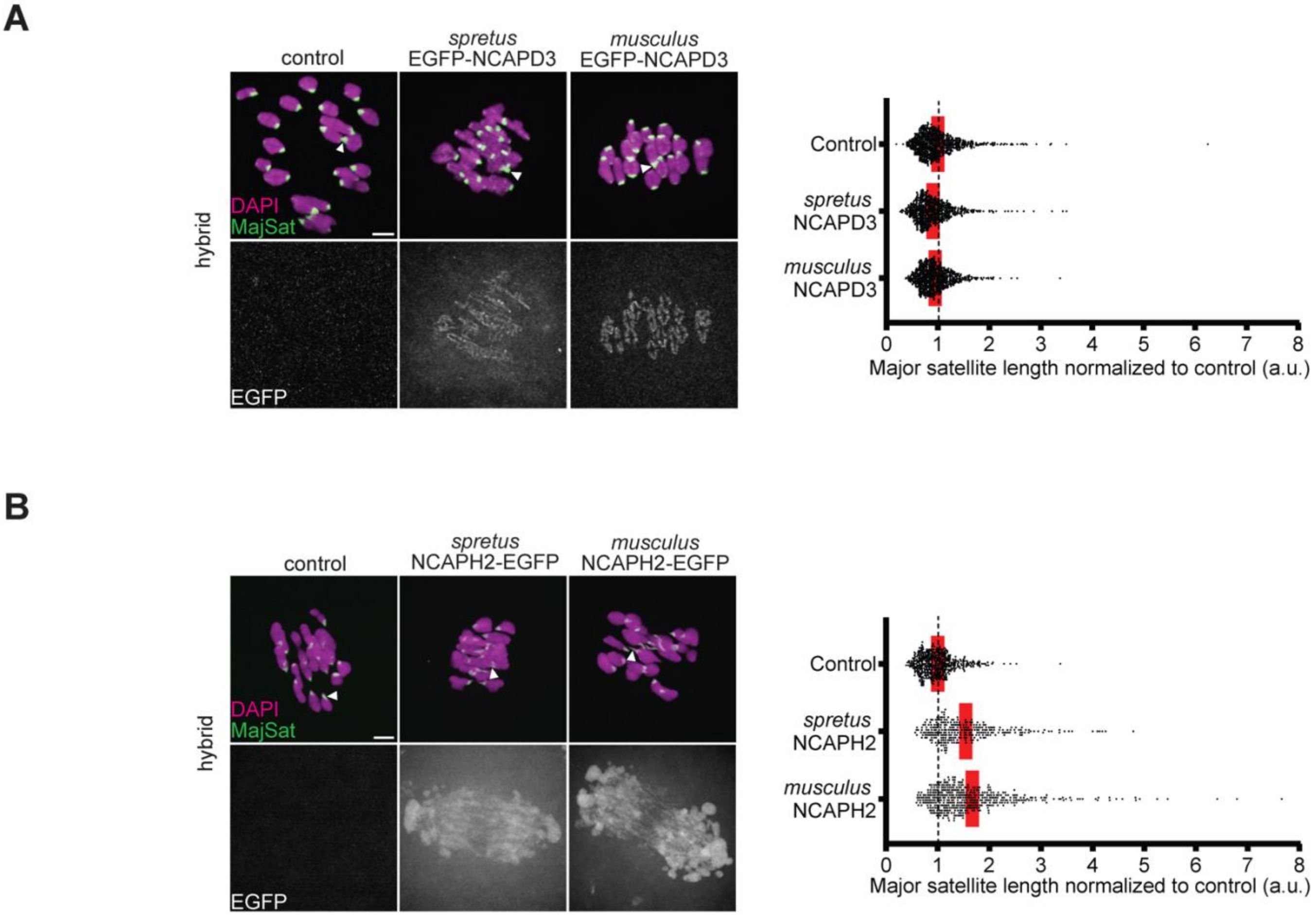


**Fig. S3.**

**Overexpressing NCAPD3 or NCAPH2 did not rescue centromere stretching.** Hybrid oocytes expressing $^{\text{EGFP-}}$NCAPD3 (**A**) or NCAPH2$^{\text{-EGFP}}$ (**B**) derived from *musculus* or *spretus* were fixed at prometaphase I and stained for TOP2A (a major satellite marker, see fig. S7B) and EGFP. Arrowheads indicate stretched centromeres. The length of major satellites was quantified; each dot in the graph represents a single centromere (n > 464 centromeres for each condition). NCAPH2$^{\text{-EGFP}}$ was not able to localize on the chromosome probably because of the competition with endogenous NCAPH2 (*21*). The enhanced centromere stretching upon overexpressing NCAPH2$^{\text{-EGFP}}$ implies that NCAPH2$^{\text{-EGFP}}$ sequesters other condensin subunits in the cytoplasm, reducing the functional condensin II pool. Images are maximum intensity z projections to show all chromosomes; red line, mean; *$P$ <0.05; scale bars: 5 µm.

NCAPG2 alignment

```
human           MEKRETFVQAVSKELVGEFLQFVQLDKEASDPFSLNELLDELSRKQKEELWQRLKNLLTD    60
musculus        MEKREAFIQAVSKELVEEFLQFLQLDKDSSNPFSLSELLDELSRKQKEELWQRLKDLLTE    60
spretus         MEKREAFIQAVSKELVEEFLQFLQLDKDSSNPFSLSELLDELSRKQKEELWQRLKDLLTE    60
                *****:*:******** *****:****::*:****.*******************:***:

human           VLLESPVDGWQVVEAQGEDNMETEHGSKMRKSIEIIYAITSVILASVSVINESENYEALL    120
musculus        TLLESPVDRWQTVEVEGADDMESEHSPKMRKSIKIICAIVTVILASVSIINEHENYGALL    120
spretus         TLLESPVDRWQTVEVEGADDMESEHSPKMRKSIKIICAIVTVILASVSIINEHENYGALL    120
                .******* **.**.:* *:**:**. ******:** **.:*******:*** *** ***

human           ECVIILNGILYALPESERKLQSSIQDLCVTWWEKGLPAKEDTGKTAFVMLLRRSLETKTG    180
musculus        ECAVILNGILYALPESEQKLQNSIQDLCVKWWERGLPAKEDMGKTAFIMLLRRSLETKSG    180
spretus         ECAVILNGILYALPESEQKLQNSIQDLCVKWWERGLPAKEDMGKTAFIMLLRRSLETKSG    180
                **.:*************:***.*******.***:******* *****:**********:*

human           ADVCRLWRIHQALYCFDYDLEESGEIKDMLLECFININYIKKEEGRRFLSCLFNWNINFI    240
musculus        ADVCRLWRIHQALYCFDYDWEESREIKDMLLECFINVNYIKKEEGRRFLSFLFSWNVDFI    240
spretus         ADVCRLWRIHQALYCFDYDWEESREIKDMLLECFINVNYIKKEEGRRFLSFLFSWNVDFI    240
                ******************* *** ************:************* **.**::**

human           KMIHGTIKNQLQGLQKSLMVYIAEIYFRAWKKASGKILEAIENDCIQDFMFHGIHLPRRS    300
musculus        KMIHETIKNQLAGLQKSLMVHIAEIYFRAWKKASGKMLETIEYDCIQDFMFHGIHLLRRS    300
spretus         KMIHETIKNQLAGLQKSLMVHIAEIYFRAWKKASGKMLETIEYDCIQDFMFHGIHLLRRS    300
                **** ****** ********:***************:**:** ************* ***

human           PVHSKVREVLSYFHHQKKVRQGVEEMLYRLYKPILWRGLKARNSEVRSNAALLFVEAFPI    360
musculus        PVHSKVREVLSYFHQ-QKVRQGVEEMLYRLYKPILWRGLKARNSEVRSNAALLFVEAFPI    359
spretus         PVHSKVREVLSYFHQ-QKVRQGVEEMLYRLYKPILWRGLKARNSEVRSNAALLFVEAFPI    359
                **************: :*******************************************

human           RDPNLHAIEMDSEIQKQFEELYSLLEDPYPMVRSTGILGVCKITSKYWEMMPPTILIDLL    420
musculus        RDPNFTATEMDNEIQKQFEELYNLIEDPYPRVRSTGILGVCKISSKYWEMMPPNILVDFL    419
spretus         RDPNFIATEMDSEIQKQFEELYNLIEDPYPRVRSTGILGVCKISSKYWEMMPPNILVDFL    419
                ****: * ***.**********.*:***** ************:*********.**:*:*

human           KKVTGELAFDTSSADVRCSVFKCLPMILDNKLSHPLLEQLLPALRYSLHDNSEKVRVAFV    480
musculus        KKVTGELAFDISSADVRCSVFKCLPIILDNKLSHPLLEQLLPTLRYSLHDNSEKVRVAFV    479
spretus         KKVTGELAFDISSADVRCSVFKCLPIILDNKLSHPLLEQLLPTLRYSLHDNSEKVRVAFV    479
                ********** **************:****************:*****************

human           DMLLKIKAVRAAKFWKICPMEHILVRLETDSRPVSRRLVSLIFNSFLPVNQPEEVWCERC    540
musculus        DLLLKIKAVRAAKFWKICPMEDILVRLEMDSRPVSRRLVSLIFNSFLPVNQPEEVWCERC    539
spretus         DLLLKIKAVRAAKFWKICPMEDILVRLEMDSRPVSRRLVSLIFNSFLPVNQPEEVWCERC    539
                *:*******************.****** *******************************

human           VTLVQMNHAAARRFYQYAHEHTACTNIAKLIHVIRHCLNACIQRAVREPPEDEEEEDGRE    600
musculus        VTLIQMNRAAARRFYQYAHEHTASTNIAKLIHVIRHCLNACIQRTLREGSEAH---KECE    596
spretus         VTLIQMNRAAARRFYQYAHEHTASTNIAKLIHVIRHCLNACIQRTLREGPEAH---KECE    596
                ***:***:***************.********************::**  * .   .  *

human           KENVTVLDKTLSVNDVACMAGLLEIIVILWKSIDRSMENNKEAKLYTINKFASVLPEYLK    660
musculus        KENASVLDKTLSVNDTASMAGLLEIIVILWKNIHRSLENNKEAKIYTINKFAAVLPEYLK    656
spretus         KENASVLDKTLSVNDTASMAGLLEIIVILWKNIHRSLENNKEAKIYTINKFAAVLPEYLK    656
                ***.:**********.*.*************.*.**:*******:*******:*******

human           VFKDDRCKIPLFMLMSFMPASAVPPFSCGVISTLRSREEGAVDKSYCTLLDCLCSWGQVG    720
musculus        VFKDERCKIPLFMLMSFFPASAVPVFSCGVISVLRNQE-SVTGRSYCTLLDCLCSWGQVG    715
spretus         VFKDERCKIPLFMLMSFLPASAVPVFSCGVISVLRNQE-SVTGRSYCTLLDCLCSWGQVG    715
                ****:************:****** *******.**.:* ....:****************
```

```
human        HILELVDNWLPTEHAQAKSNTASKGRVQIHDTRPVKPELALVYIEYLLTHPKNRECLLSA    780
musculus     HVLELIVDWLPTVPPQAKSNLASKRKVEINDTCSVKPELALLYMEYLLTHPKNRECLLSV    775
spretus      HVLELIVDWLPTVPPQAKSNLASKRKVEINDTCSVKPELALLYMEYLLTHPKNRECLLSV    775
             *:***: :****   ***** *** :*:*:**  *******:*:***************.

human        PRKKLNHLLKALETSKADLESLLQTPGGKPRGFSEAAAPRAFGLHCRLSIHLQHKFCSEG    840
musculus     PQKKLNQLLKALEGSKAELESFLQSPSGNPLNFNKATALHAFGLYCRMSVHLQYKFCSEE    835
spretus      PQKKLNQLLKALEGSKAELESFLQSPSENPLNFNKATALHAFGLYCRMSVHLQYKFCSEE    835
             *:****:****** ***:***:**:*. :* .*.:*:* :****:**:*:***:*****

human        KVYLSMLEDTGFWLESKILSFIQDQEEDYLKLHRVIYQQIIQTYLTVCKDVVMVGLGDHQ    900
musculus     KIHLSILDDTGSWLENKVLPLLEDQEEEYLKLRKDVYQQIIQTYLAVCKDVVMVGLGDPK    895
spretus      KIHLSILDDTGSWLENKVLPLLEDQEEEYLKLRKDVYQQIIQTYLAVCKDVVMVGLGEPK    895
             *::**:*:*** ***.*:* :::****:****:: :*********:***********: :

human        FQMQLLQRSLGIMQTVKGFFYVSLLLDILKEITGSSLIQKTDSDEEVAMLLDTVQKVFQK    960
musculus     FQMQLLQRSFGIMKTVKGFFYVSLLLGILKEIAGNTIIHKTDSDEKVTVLFDLVQEVFQK    955
spretus      FQMQLLQRSFGIMKTVKGFFYVSLLLGILKEIAGNTIIHKTDSDEKVTVLFDLVQEVFQK    955
             *********:***:************.*****:*.::*:******:*::*:* **:****

human        MLECIARSFRKQPEEGLRLLYSVQRPLHEFITAVQSRHTDTPVHRGVLSTLIAGPVVEIS    1020
musculus     MLECIACIFRKQPEESLPLFHSVQTPLHEFITTIQSWHKDTAVHHAVLSTLIAAPVVEIS    1015
spretus      MLECIACIFRKQPEESLPLFHSVQTPLHEFITTIQSWHKDTAVHHAVLSTLIAAPVVEIS    1015
             ******  *******.* *::*** *******::** *.** **:.*******.******

human        HQLRKVSDVEELTPPEHLSDLPPFSRCLIGIIIKSSNVVRSFLDELKACVASNDIEGIVC    1080
musculus     HQLQKVSDIEELTSPQCLHDLPPFSRCLVGVIMKSSDVVRSFVDELKACVTSGDVEGIVC    1075
spretus      HQLQKVSDIEELTSPQCLHDLPPFSRCLVGVIMKSSDVVRSFLDELKACVTSGDVEGIVC    1075
             ***:****:**** *: * *********:*:*:***:*****:*******:*.*:*****

human        LTAAVHIILVINAGKHKSSKVREVAATVHRKLKTFMEITLEEDSIERYEDL--LCCPGWA    1138
musculus     LTAVLHIILVINKGKHISAKVKEVATAVYRKLKTFMEITLEEDSLERFLYESSMRTLGEF    1135
spretus      LTAVLHIILVINKGKHISAKVKEVAAAVYRKLKTFMEITLEEDSLERFLYESSMRTLGEF    1135
             ***.:******* *** *:**:***::*:***************:**:     :   *

human        LTPGLLDSIYPSASVPIG    1156
musculus     LNP---------------    1138
spretus      LNP---------------    1138
             *.*
```

**Fig. S4.**

**Amino acid sequence alignment of NCAPG2.** Amino acid sequence of NCAPG2 from *Homo sapiens*, *Mus musculus domesticus*, and *Mus spretus* were aligned. Residues diverged between *musculus* and *spretus* were highlighted in yellow. Although there are several residues diverged between *musculus* and *spretus* NCAPG2, these differences are probably not part of the hybrid incompatibility, because both of them efficiently rescued centromere stretching when overexpressed in hybrid oocytes (Fig. 2C). NCAPG2 antibodies used in this study (Bioss bs-7721R and Bethyl A300-605A) were raised against human NCAPG2 fragments. Antigen sequences used for the antibody productions were highlighted in light green for the Bioss antibody and light blue for the Bethyl antibody. Since the sequences are basically conserved between *musculus* and *spretus* for the region corresponding to the antigen sequences, it is unlikely that the distinct NCAPG2 staining patterns among *musculus*, hybrid, and *spretus* oocytes are due to the differential efficiency in antibody recognition (Fig. 3A and fig. S5B).

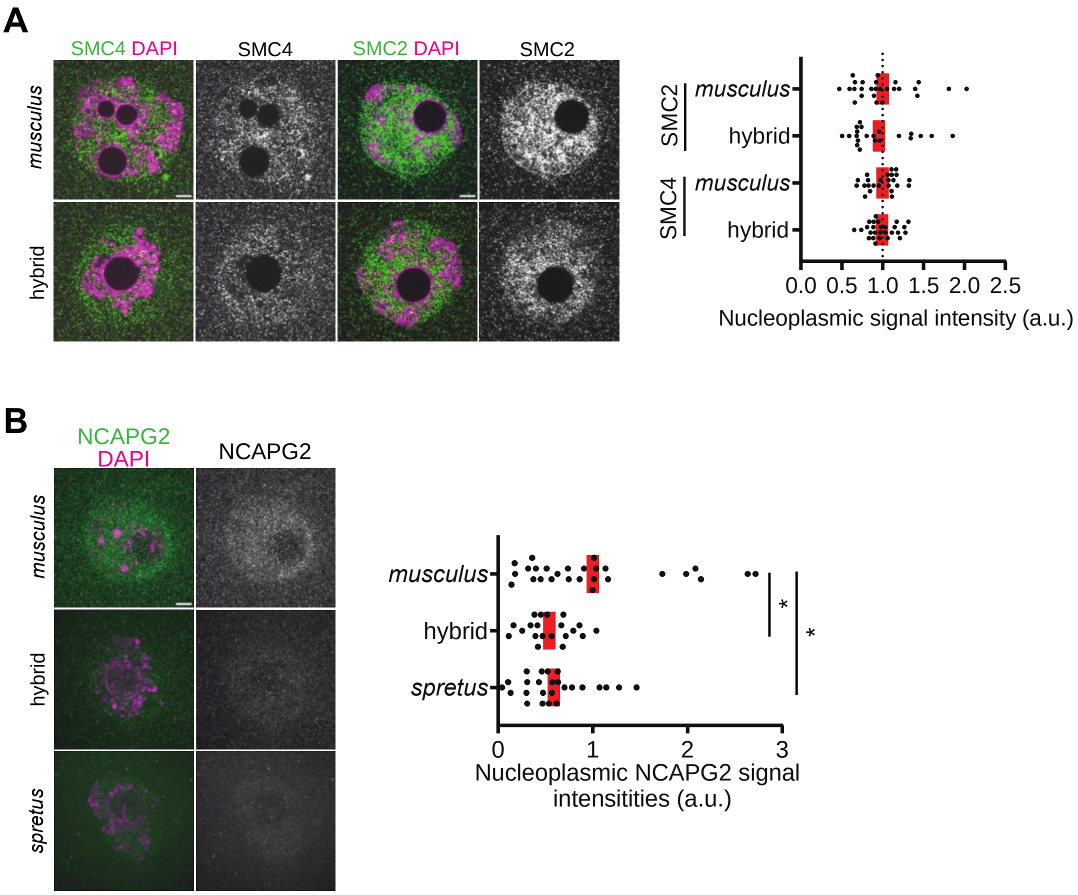


**Fig. S5.**
**Nuclear enrichment of condensin II subunits in late prophase I.** (**A**) *musculus* and hybrid oocytes were fixed at prophase I and stained for SMC2 and SMC4. The graph shows the quantification of SMC2 and SMC4 intensities in the nucleus (n > 20 oocytes for each condition). (**B**) *musculus*, hybrid, and *spretus* oocytes were fixed at prophase I and stained for NCAPG2 (Bethyl A300-605A). The graph shows the quantification of NCAPG2 intensities in the nucleus, which shows a similar trend with Fig. 3A where a different NCAPG2 antibody (Bioss bs-7721R) was used (n > 25 oocytes for each condition). Images are optical slices to show the nucleus; red line, mean; **P* <0.05; scale bars: 5 $\mu$m.

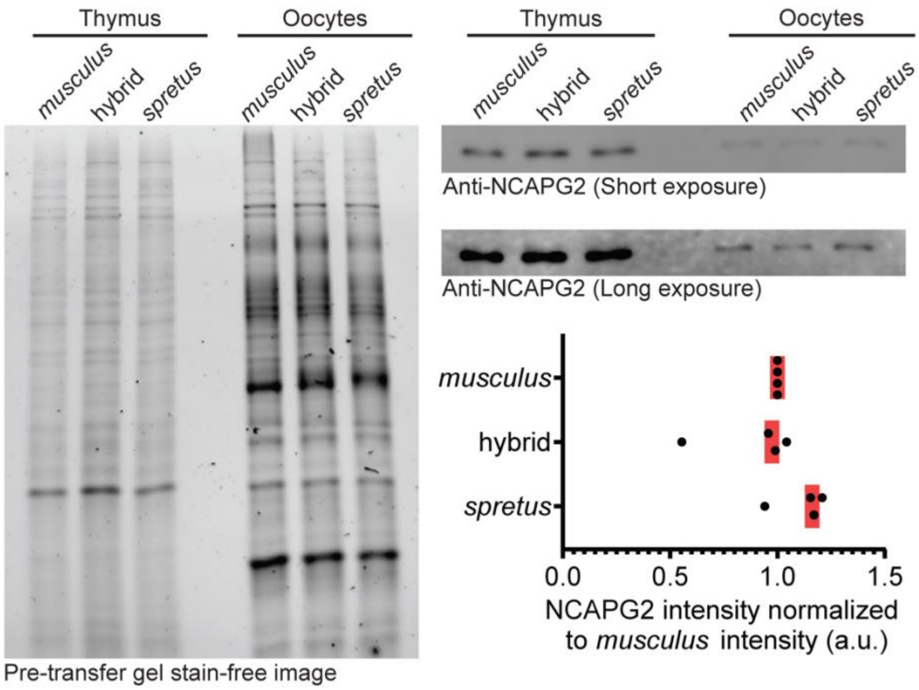


**Fig. S6.**

**Total NCAPG2 protein levels are equivalent among *musculus*, *spretus*, and hybrid oocytes.**
Westernblot of NCAPG2 using thymus tissues and oocyte samples. As a loading control, stain-free gel image was shown to indicate the total protein amount loaded to each lane. Oocyte collection and westernblot were repeated four times to quantify the total NCAPG2 protein level in each genotype.

256

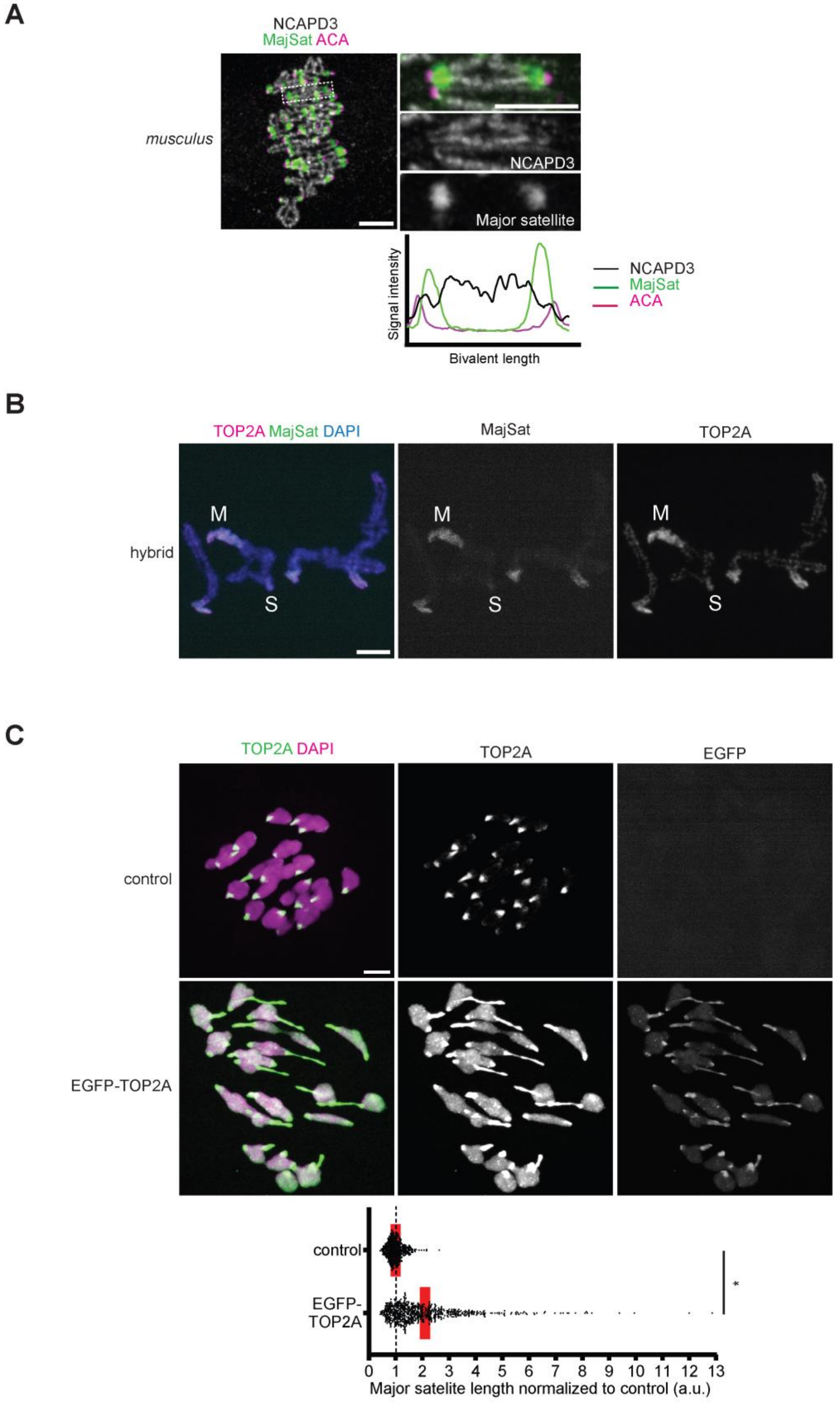

A
NCAPD3
MajSat ACA
musculus
NCAPD3
Major satellite
Signal intensity
Bivalent length
NCAPD3
MajSat
ACA
B
TOP2A MajSat DAPI
MajSat
TOP2A
hybrid
M
S
C
TOP2A DAPI
TOP2A
EGFP
control
EGFP-TOP2A
control
EGFP-TOP2A
*
0 1 2 3 4 5 6 7 8 9 10 11 12 13
Major satelite length normalized to control (a.u.)

**Fig. S7.**

**High TOP2A and low condensin II abundance at major satellites.** (**A**) *musculus* oocytes expressing MajSat were fixed at metaphase I and stained for NCAPD3 (antibody from the Hirano lab). The graph is line scans of MajSat (green), centromere (magenta), and NCAPD3 (black) intensities across the chromosome length. (**B**) Chromosome spreads were performed at metaphase I from hybrid oocytes expressing MajSat and stained for TOP2A. “M” and “S” indicate *musculus* and *spretus* centromeres, respectively. *musculus* centromeres with major satellites highly enrich TOP2A while *spretus* centromeres enrich significantly less. (**C**) Hybrid oocytes expressing $^{\text{EGFP-}}$TOP2A were fixed at prometaphase I and stained for TOP2A and EGFP. The major satellite length was quantified (n > 543 centromeres for each condition). Overexpressing TOP2A enhanced the centromere stretching phenotype rather than rescuing it. Images are maximum intensity z projections to show all chromosomes or optical slices magnified to show individual chromosomes; red line, mean; $^*P < 0.05$; scale bars: 5 $\mu$m.

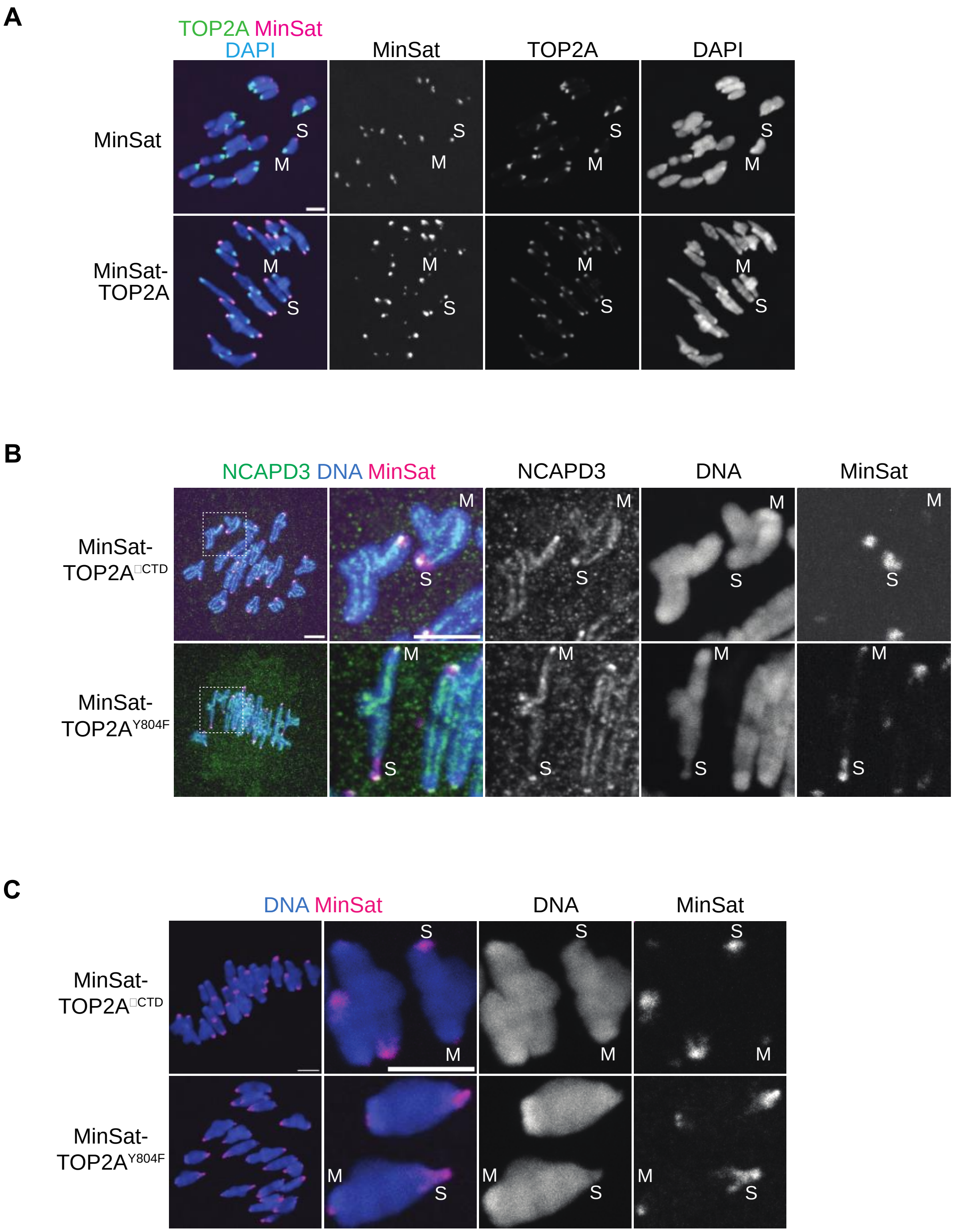


**Fig. S8.**

**TOP2A's catalytic activity and its guiding function to localize to the chromosome axis are critical to reduce condensin II abundance at centromeres.** (**A**) Hybrid oocytes expressing MinSat or MinSat-TOP2A were fixed at prometaphase I and stained for TOP2A. TOP2A abundance recruited to *spretus* centromeres by this targeting strategy was equivalent to that on major satellites. (**B**, **C**) Hybrid oocytes expressing MinSat-TOP2A$^{Y804F}$ or MinSat-TOP2A$^{\Delta CTD}$ were fixed either at metaphase I and stained for NCAPD3 (B) or fixed at prometaphase I and counterstained with DAPI (C). MinSat-TOP2A$^{Y804F}$ and MinSat-TOP2A$^{\Delta CTD}$ are less efficient in reducing condensin II abundance and inducing centromere stretching compared to MinSat-TOP2A (wild-type), indicating that both TOP2A's catalytic activity and its guiding function to localize to the chromosome axis are important to reduce condensin II abundance at centromeres (*30*). "M" and "S" indicate *musculus* and *spretus* centromeres, respectively. Images are maximum intensity z projection to show all chromosomes (left) or optical slices magnified to show individual chromosomes (right). Scale bar: 5 $\mu$m.
